# How alginate lyase produces quasi-monodisperse oligosaccharides: a normal mode analysis-based docking and molecular dynamics simulation study

**DOI:** 10.1101/2021.07.25.452152

**Authors:** Hengyue Xu, Qi Gao, Dengming Ming

## Abstract

Polysaccharide degradation products are widely used in medicine, health food, textile and other industries. The preparation of monosaccharides by enzymatic degradation is a key technology in bio industrial production. Unfortunately, most of the known digested products are complex oligosaccharide mixtures, which limit their industrial processing and application. In this study, we explored a docking technique based on normal mode analysis to examine the possible cleavage mechanism of an alginate lyase (AlyB) from *Birio Splendidus*, which contains the catalytic domain of polysaccharide lyase family 7 (PL7) and a CBM32 sugar binding module, and was observed to produce trisaccharide products with quasi-monosaccharide distribution. We compared the molecular interactions of the enzyme with the natural alginates, the polyMG whose products has the quasi-monodisperse distribution of tri-saccharide and two synthetic polysaccharides, the polyM and polyG whose products has a wider distribution of oligosaccharides. Our calculations quantitatively show that there are a series of deterministic conformational changes in the catalytic pocket, which control the specificity of the substrate; at the same time, it determines the uniformity of the final product together with the spatial position of the key catalytic sites. The dynamic simulations revealed that CBM domain plays a key role in assisting the release of tri-saccharides. Our data highlights the important role of enzyme flexibility in determining product uniformity, which may provide new insight into design of enzymes for the production of high-value mono-distributed oligosaccharides.

## Introduction

As an effort to meet the fossil fuel depletion and the climate change problem, scientists had been looking for alternative green energy sources, among which macroalgae is considered to be a typical green energy because it can grow with a large amount of biomass without consuming a lot of land. Macroalgae, such as *Laminaria, macrocystis, Ascophyllum*,etc., are rich in polysaccharides that has the major form of sodium alginate in the algal cell wall. Alginates are polysaccharides that are comprised of *α*-L-Guluronate (G) and *β* −D-mannuronate(M), which are linked to form either homo-polymeric blocks (poly-G or poly-M) or alternating/random heteropolymeric blocks (poly-MG or poly-GM) of varying sizes via a 1,4-glycosidic linkage. The flexible arrangement and combination of M and G units endows alginates with a broad-spectrum of structures and functions.

Alginates are degraded by the so-called alginate lyases that catalyze the degradation of by the mechanism of β-elimination of glycosidic bonds, producing unsaturated oligosaccharides with an uronic acid moiety at the non-reducing terminal [1]. Over the past decades, thousands of alginate lyases have been identified as capable of degrading and modifying the fine structure of alginates to produce a variety of alginate oligosaccharides with potential antioxidant, antipathogenic, anti-inflammatory and other bioactive properties as well as the biofuel sources[2, 3]. Alginate lyases are currently classified into 12 polysaccharides lyases (PL) families (PL5, PL6, PL7, PL14, PL15, PL17, PL18, PL31, PL32, PL34, PL36 and PL39) in CAZy database[4], according to their sequence similarity, domain functionality and substrate specificity [5-7]. The diversity of structure and morphology of alginate determines the existence of a variety of alginate-lyases with similar function but different structure and substrate specificity, which provides a good example for the study of the function and structure specificity of enzymes[8].

Most products of alginate lyases are mixtures of oligosaccharides and the separation of these mixtures is usually very difficult or costly, limiting their application[5, 9, 10]. Many alginate lyases have monosaccharides as their main product, while others have a product of the mixtures of disaccharides to tetra-saccharides[5]. Among the various alginate lyases, those that produce quasi-monodisperse oligosaccharides with specific types of alginate substrates are of particular interest, due to their potential bioengineering and medical applications. For example, Lyu etc. reported the structure of a multidomain alginate lyase isolated from *Vibrio spendidus* OU02, named AlyB, which is composed of a PL catalytic domain and a carbohydrate binding modules (CBM32 domain) [11]. In the structure, the two domains are separated without any direct contact, but kept connected with a noncanonical alpha helix linker. Their data suggested that the presence of the rigid linker helix and the CBM32 domain makes the enzyme product predominantly trisaccharide. Recently, we also screened an alginate lyase from *Vibrio spendidus* W13, named ALY7A, which is also a member of the PL7 family[12]. Aly7A is a bifunctional alginate lyase that shows enzymatic activities against PolyMG or PolyGM at an optimal temperature of 30°C and pH of 7. Interestingly, Aly7A exhibits a specific degradation pattern that specifically produces tri-saccharides as the main product of three different substrate types[13]. It also converts tetra-saccharides to monosaccharides and tri-saccharides, and penta-saccharides to disaccharides and tri-saccharides. A structural modeling revealed that Aly7A has 70% structural similarity of AlyB, with both full-length enzymes having similar domain arrangement and connection manner. The experimental data suggests that CBM and α-helical linker jointly control the release of the products of three saccharide units, so that the products are conducive to the mono distribution of tri-saccharides. However, the underlying catalysis and release mechanism that determines the quasi mono-distribution of polysaccharide degradation are still unclear, which hinders the development of enzymatic degradation of alginates and the relevant applications in bioengineering and pharmaceutical industries.

In this paper, the detailed interaction between alginate and the selected alginate lyase was studied by computational methods, with special attention to the details of the relative motion of the domain, the mechanism of action of alginate at the catalytic site, and their correlation with the production of quasi single release products.

In this paper, computational methods are used to study the detailed interactions between alginates and the selected alginate lyases, with special attention to the relative motions of the domains, the possible actions at the catalytic sites, and their correlation with the quasi mono-distribution of the released products. We explored a series of configurations that the enzyme can sample in docking with the 3 typical forms of alginates (polyM, polyG, and polyMG), and characterized molecular interactions and binding energies between the enzyme and the substrates. Further, molecular dynamic simulations were performed to explore the interactions between the CBM domain and the product.

## METHODS

### Normal mode analysis of the molecular structure of AlyB

The 3D coordinates of AlyB is taken from the crystal structure (PDB[14] entry code 5ZU5[11]). The alginate lyase of vibrio spendidus (AlyB) comprises of two large domains: the carbohydrate binding modules CBM32 and a catalytic domain where polysaccharide is cleaved. In the structure, the two domains are linked through noncanonical rigid alpha helix that contains 26 amino acids. The complex structure leaves a lot of room for the relative movement of the two domains. To demonstrate these possible motions, a normal mode analysis (NMA) method was used[15]. NMA provides a shortcut to identify important allosteric conformation changes accessible to biostructures without time-consuming molecular dynamic simulations, especially when protein functional conformations are not far from their equilibrium structures[15, 16]. As a simplified version of the conventional all-atom NMA, here we used an anisotropic version of elastic network model (ANM)[17, 18] to derive the direction of movement of all the heavy atoms of the structure. This model had been widely used to model key conformation changes of a variety of biostructures[16, 19, 20] as well as multi-scaled simulations[21]. In ANM[17, 20], every protein atom is constrained not to move away from its surrounding atoms within a cutoff distance of 7Å through an empirical harmonic potential ***U***. A diagonalization of the Hessian matrix, a matrix whose elements are the mass-weighted second derivatives of the potential ***U*** with respect to atoms’ spatial coordinates, gives a series of eigenvectors **V**’s, or normal modes, each of them specifies the orientation and amplitude for each atom to move. The normal mode defines how the structure changes its shape near the equilibrium conformation. Each eigenvector or normal mode goes with a scalar value, called eigenvalue, which is square of the frequency *f* – a value of time factor e^*iωt*^, where ;*ω* = 2*πf*, that measures how fast the structure adjusts its shape around an equilibrium conformation. The mode is simply demonstrated by arrows applied to C-alpha atoms in the structure whose orientation and amplitude was given **V** cos(*θ*).

### Docking AlyB with alginate

We first constructed the AlyB and alginate substrate complex by molecular docking simulation. The molecular docking studies were performed using the Schrödinger suite (version 2017). The crystal structure of AlyB (PDB code 5ZU5) was prepared using the “Protein Preparation Wizard” module in Schrodinger Maestro (version 11.0) with the default protein parameters. Hydrogen atoms were added, hydrogen bonds were optimized, partial charges for all atoms were assigned, and hydrogen atoms were energy-minimized using the OPLS-3e force field. Finally, all crystal water was Removed the receptor grid with Schrodinger Glide, the Y466 center of mass was defined as the grid center; the size of the site was set to 36Å, which is adequate enough to accommodate the hexamer polysaccharide. The hexamer polyM was built using Schrodinger Maestro (version 11.0), the protonation state was calculated using the Schrodinger Epik, and 20 conformations of the hexamer were generated using Schrodinger ConfGen. All conformers were docked to the prepared protein structure using Glide (docking parameters: standard precision, dock flexibly, All constraints in grid were activated).

### Molecular dynamics simulation

To understanding the function of CBM domain Simulations were performed by GROMACS[22] software. Charmm36 force field was used to generate the topology file of the protein, choosing TIP3P water model. A topology file of 3mg / 9mg was generated using the CGenFF[23, 24] server, and the final complex was constructed. The initial structure of each system was solvated in a cubic box with a box length of 119.98Å. An appropriate amount of sodium counterion was added to neutralize each system so that it remained electroneutral. Prior to equilibration, to constrain the ligand, we generated a position restrained topology for it. To minimize the energy, the system was optimized using the steepest descent minimization. Subsequently, the protein was subjected to a position restraint, after which each system was slowly heated to 300 K with a constant volume and equilibrated. Finally, a MD equilibration simulation of 200 ns was performed. Short range van der Waals cutoff was calculated using the Verlet algorithm and long range electrostatics was calculated using the particle mesh Ewald method (PME)[25]. The time step was set to 2 fs, and trajectories were collected at 10 ps intervals. The trajectories were analyzed using the gmx built-in programs -rms, -gyrate, -rmsf.

## Results and Discussion

### The polysaccharide-cleavage chemical mechanism of AlyB

To explore the chemical mechanism of polysaccharide-cleavage in AlyB, the molecular docking simulation was used to study the AlyB and Glycan substrate complexes. The molecular structure, binding energy, bond length, bond Angle and distance of lyase-glycan complex were obtained. This information was analyzed in detail and the mechanism of polysaccharide cleavaging by alginate lyase was chemically summarized.

Figure 1a) showed the protein structure where Tyr466 and Trp129 were identified as key catalytic residues. Tyr466 is an important site for the PD-7 domain to perform catalytic functions, while Trp129 is an important site for the CBM32 domain to bind product tri-saccharides. From the charge surface diagram of amino acids in Figure 1b), it can be observed that the protein surface of AlyB contains a large number of charged residues. This complex charge environment creates conditions for the binding of the glycan chain of AlyB and the cleavage chemical reaction of polysaccharides.

**Figure 1.**
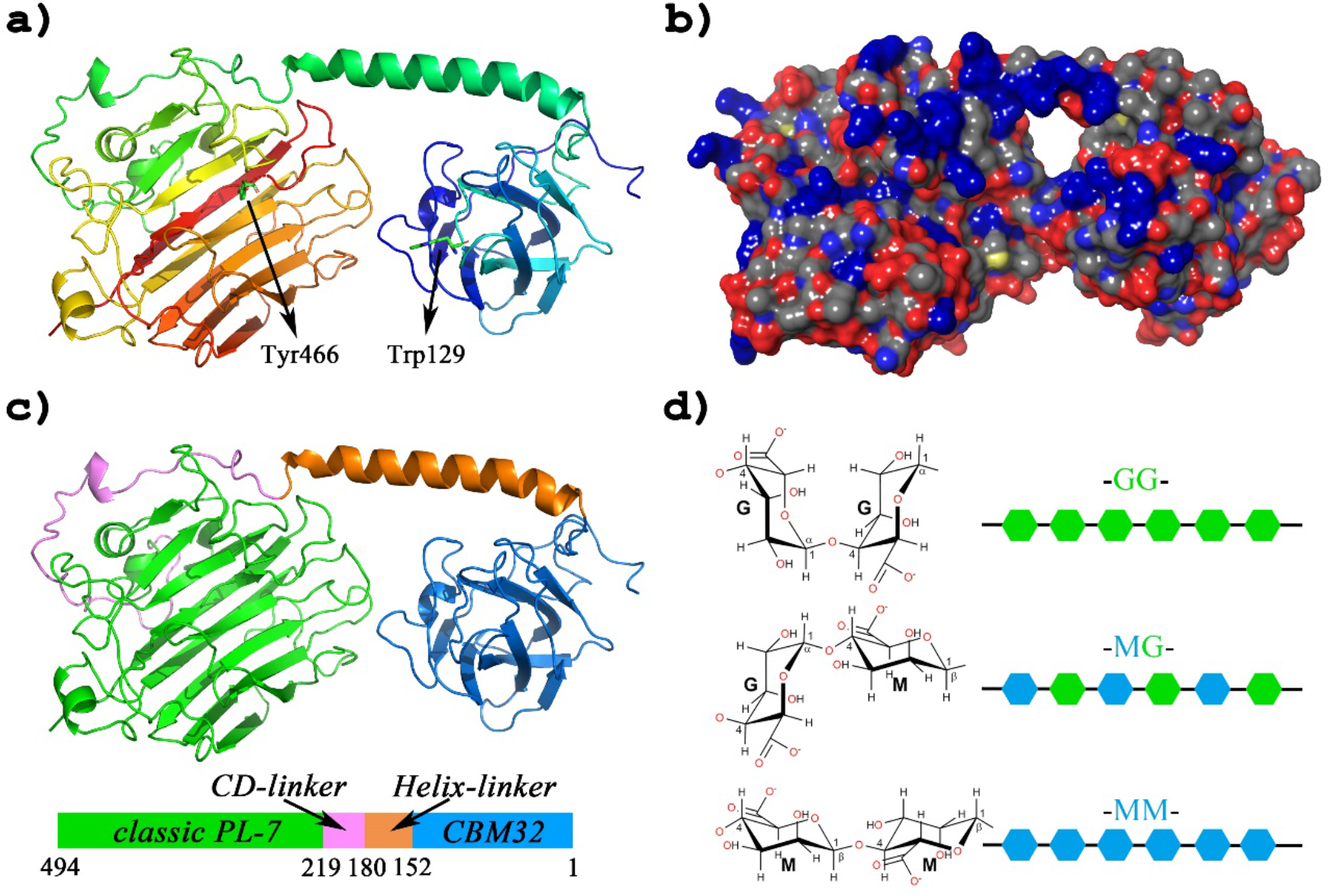
(a). The structure of AlyB (PDB code:5ZU5) is shown as cartoon (in ranbow spectrum mode). Key residues have been marked with black arrows. (b). The molecular surface of AlyB built by pymol (colored by residue charge). (c). The protein is divided into four components, PL-7 (in green), CD-linker (in violet), Helix-linker (in orange), and CBM32 (in blue). (d). Three configurations alginate (green hexagon represent G and blue hexagon represent M), natural alginate polyM (-MG-) and two artificial syntheses alginate polyM (-GG-) and (-MM-).

In Figure 1c), AlyB is divided into four components, PL-7, CD-linker, Helix-linker, and CBM32. In previous reports, PL-7 has been proved to be a domain with the activity of polysaccharide cleavage reaction, while CBM32 is believed to have an accurate cleavage effect on the lytic activity of AlyB polysaccharides, but it has little effect on the catalytic reaction itself[11].

Helix-linker is a domain linker that is more rigid than random coils, and its length will affect the interaction between the two domains PL-7 and CBM32, which may affect the molecular weight distribution of the product. In the previous literature, it was surprising that the effect of CD-linker was essential for the cleavaging chemical activity of polysaccharides. The loss of Helix-linker and CBM32 affected only the accuracy of the cleavage reaction and had little effect on the reactivity, while the loss of CD-linker resulted in the complete disappearance of catalytic reactivity.

As for ligand molecules, we use the nine-glycan of alginate to approximate the macromolecular chain of alginate. Alginate can be divided into artificial alginate and natural alginate. Artificial alginate is a single β-D-mannuronic acid (M) or α-L-guluronic acid (G) polymer, while natural alginate is an alternate polymer with M and G arranged alternately. As shown in Figure 1d), there are three configurations of the alginate molecular chain, namely -GG-, -MG-and -MM-. The different arrangement of M and G will inevitably lead to the different physical and chemical properties of the three glycan chains, such as molecular geometry, surface area and flexibility, etc., which will also lead to the different degree of binding difficulty of AlyB to different alginate.

Then we first used alginate 9-glycan for molecular docking with AlyB. Fig. S1, S2 and S3 reveal a series of conformations generated by artificial alginate 9MM, 9GG and natural alginate 9MG docking with AlyB molecule respectively. We found that the spatial distribution of a series of docking structures obtained by artificial alginate 9MM and 9GG was relatively dispersed (as shown in Figure S1 and S2).

Moreover, it can also be observed from Table 1 that the distance between Osite and Y466-OH in the 10 conformations generated by 9MG docking is relatively close, ranging from 1.96 Å to 5.10 Å. For 9MM and 9GG, the range is 2.80 Å∼17.69 Å and 4.24 Å∼14.02 Å, respectively. Most structures have a large distance and the numerical range of distance is large. Based on the above spatial structure statistics, it is revealed that the combination of artificial alginate 9MM and 9GG with alginate is not very stable, which is also the reason why the experimental hydrolysis of artificial alginate by AlyB did not yield oligosaccharides with relatively uniform molecular weight **[?]**. However, the spatial distribution of a series of complexes obtained by the docking of natural alginate 9MG with AlyB is much denser than that of artificial alginate, and the spatial positions of most conformations are not much different. In the binding energy analysis, we found that the binding energies of artificial alginate 9MM, 9GG with AlyB were -8.550 kcal/mol^-1^ and -8.487 kcal/mol^-1^, respectively. However, the binding energy of natural alginate 9MG with AlyB was - 9.890 kcal/mol^-1^, which was -1.34 kcal/mol^-1^ and -1.403 kcal/mol^-1^ lower than that of artificial alginate 9MM and 9GG, respectively. The calculated energy evidence suggests that AlyB is more likely to bind to the natural alginate 9MG. This provides a structural and energy basis for the precise cleaving of alginate by AlyB to produce triosaccharide oligosaccharides[11].

**Table 1.**
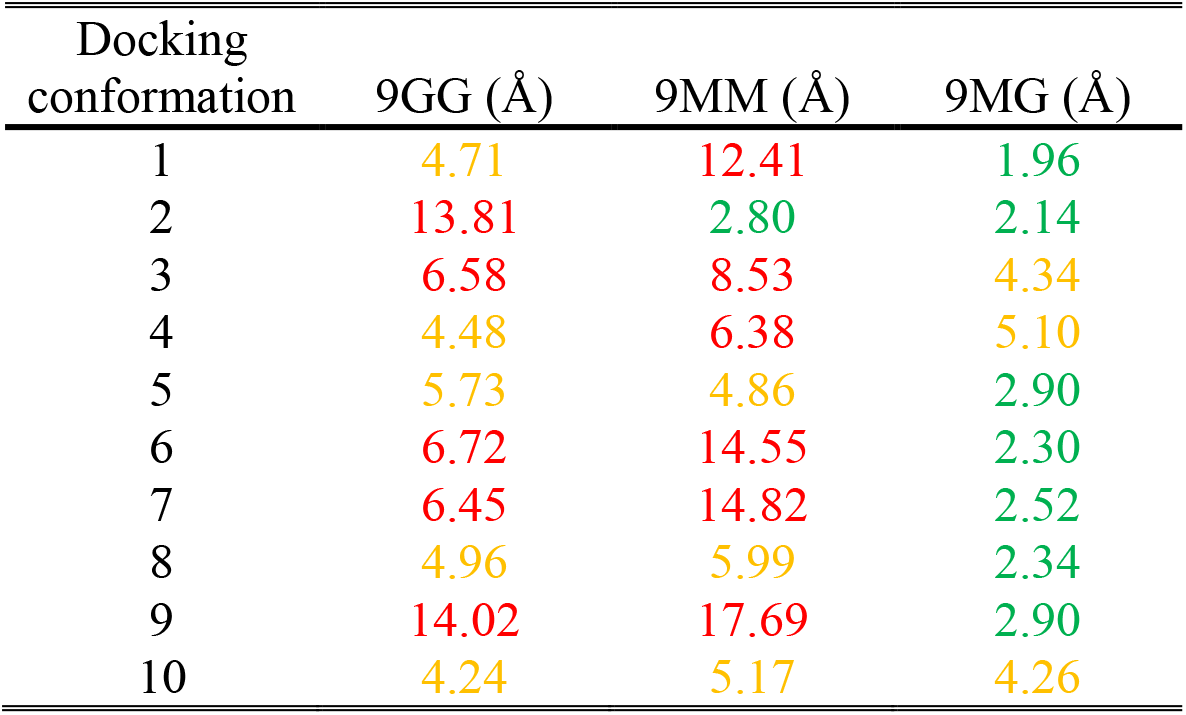
Docking binding energy from glidescore mode. In the ten conformations of alginate produced by docking, the distance between the catalytic site Y466-OH and the oxygen atom of O-glycoside between the third sugar ring and the fourth sugar ring from the right. (green: 1 Å∼4 Å, yellow: 4 Å∼6 Å, red: 6 Å∼20 Å.)

| Docking<br>conformation | 9GG (Å) | 9MM (Å) | 9MG (Å) |
| --- | --- | --- | --- |
| 1 | 4.71 | 12.41 | 1.96 |
| 2 | 13.81 | 2.80 | 2.14 |
| 3 | 6.58 | 8.53 | 4.34 |
| 4 | 4.48 | 6.38 | 5.10 |
| 5 | 5.73 | 4.86 | 2.90 |
| 6 | 6.72 | 14.55 | 2.30 |
| 7 | 6.45 | 14.82 | 2.52 |
| 8 | 4.96 | 5.99 | 2.34 |
| 9 | 14.02 | 17.69 | 2.90 |
| 10 | 4.24 | 5.17 | 4.26 |

Fig. 2a shows the molecular docking diagram of the 9MG chain of natural alginate (carbon atoms are shown in blue) bound to lyase. The catalytic site on the PL-7 domain and the key amino acid Trp129 on the CBM32 domain are marked with black arrows. We can clearly observe that 9-glycan is tightly bound to catalytic channels in AlyB, which are composed of several arcuate β-turns secondary structures. Most of the catalytic sites of the binding conformation are located between the third and fourth sugar rings, indicating that the polysaccharide cleavaging chemical reactions in natural alginate 9MG often occur between the third and fourth sugar rings, so most of the products obtained are trisaccharide oligosaccharides.

**Figure 2.**
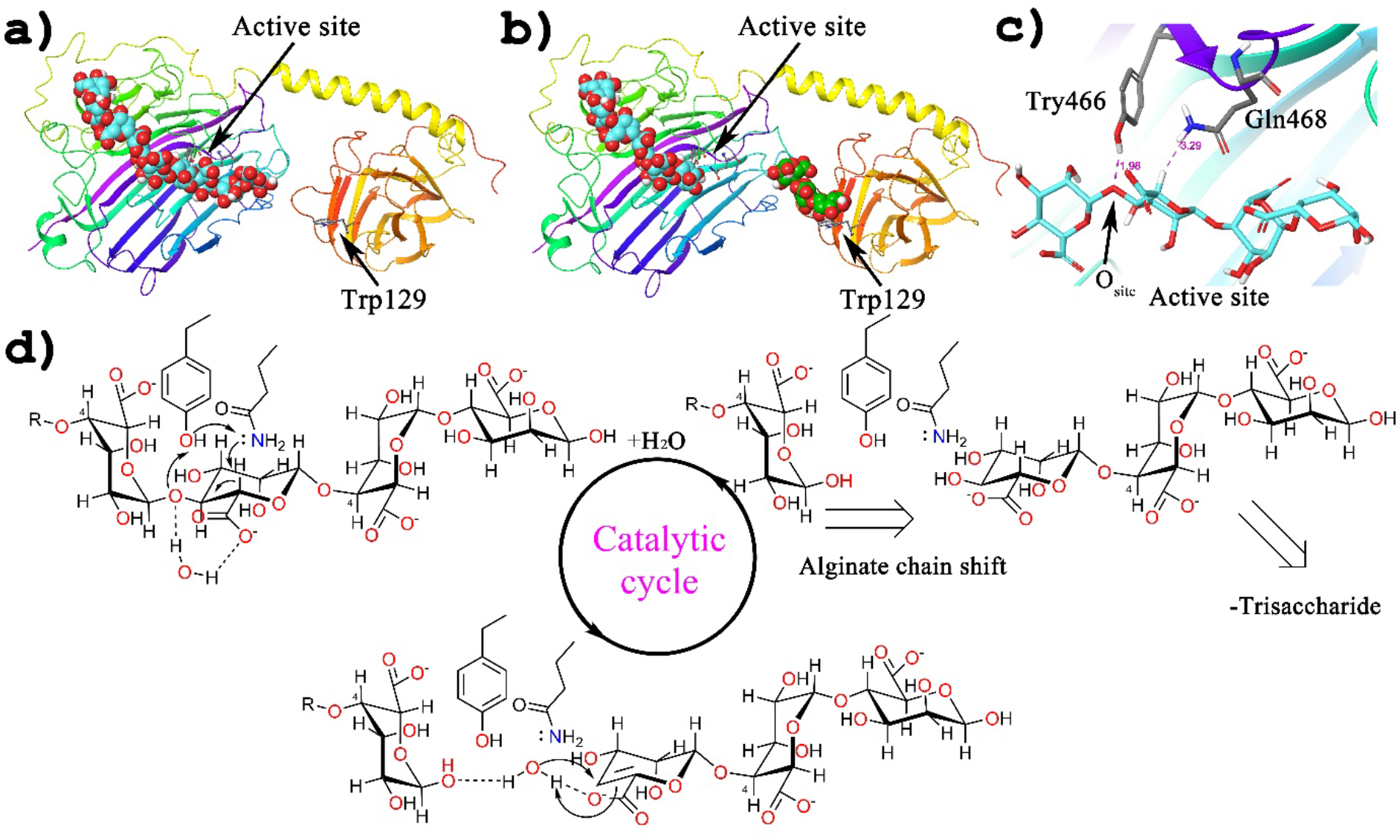
Molecular docking of alginate onto the AlyB full-length structure helped to construct a trisaccharide release model and the alginate hydrolysis chemical reaction mechanism. The annotated lengths are showed in angstroms.

The process from Figure 2a to Figure 2b is the process of cleavaging natural alginate to obtain trisaccharide. We continued to conduct molecular docking between the triosaccharide oligosaccharide generated by cleavage (carbon atoms in green) and the AlyB combined with 6-glycan, and found that the generated triosaccharide oligosaccharide would interact with Trp129 on CBM32 (Figure 2b). This indicates that CBM32 immediately binds to the product triosaccharide oligosaccharide after the cleavage of AlyB, making space for the alginate oligosaccharide chain to move and cleavage the next triosaccharide oligosaccharide in the arcuate β-turn catalytic channel. This phenomenon is completely consistent with the experimental report that CBM32 acts as the binding product triosaccharide oligosaccharide without affecting the catalytic activity[11].

Combined with the experimental reports[26], we believe that Tyr 466 and Gln 468 are likely to be directly involved in the β-elimination reaction on the third sugar ring of 9MG (Fig. 2c). We observed that the hydroxyl group on Tyr466 is only 1.96 Å away from the O-glycosides between third and fourth sugar rings in 9MG, and it is possible to form hydrogen bonds between them. The distance between the amino group in Gln468 and the acidic β-hydrogen in the third sugar ring of 9MG is 3.92 Å. Here, combining our simulation results with reported experiments, we propose a chemical reaction mechanism for cleavage alginate: The β-hydrogen on the carbon attached to the carboxylate on the six-membered sugar ring is readily dissociated and transferred to the Gln468 amino group at a distance of 3.29 Å due to the electron-withdrawing conjugation effect of the carboxylate. At the same time, the phenolic hydroxyl hydrogen of Tyr466 readily transfers to the O-glycosidic bond only 1.98 Å away from it, resulting in the breaking of the glycosidic chain and the formation of a double bond. The positive aminylium ion of Gln468 transfers hydrogen back to the phenolic hydroxyl group of Tyr466 (Fig. 2d left). Then, the surrounding solvent water molecules are added to the double bond (Fig. 2d below). The molecular chain of alginate moves to the right, back to the beginning, and performs the next cleavage reaction (Figure 2D, right). This is an insteresting catalytic cleavage cycle.

### The Normal mode based docking analysis of the catalytic mechanism of AlyB

Here, we considered the normal mode with the lowest energy thus having the longest period or smallest non-zero frequency. Figure 1(a) lists 12 snapshots of this mode, by setting the 12 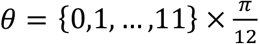. In this mode, the two domains carry out a twist motion with the linker helix as the rotation axis. As a result of the motion, the two domains continuously adjust their orientation so that they can effectively change relative distance between the two sugar-binding centers possibly up-to a large-scale while costing the smallest energy.

The 12 snapshots obtained from the Normal model were docking with natural alginate 9MG, artificial alginate 9MM and 9GG respectively, and the original crystal structure obtained from the experiment was used as the starting structure of the Normal model. In Figure 3, for the natural alginate-AlyB system (green), we observe that the docking energy of the substrate and the different conformation of AlyB is higher than that of the original crystal structure. This indicates that the original crystal structure of AlyB is the most conducive to combining with natural alginate 9MG. In addition, through normal mode analysis, we believe that AlyB has two states in the process of motion, namely Relax state (00-06) and Tense state (06-12). It is observed that the energy of AlyB is generally high in the Tense state, and alginate may be catalyzed to cleavage in the Tense state of AlyB. As for the Relax state (00-06) of AlyB, its energy fluctuates greatly, and we believe that the cleavage products triosaccharide oligosaccharides may be released in the Relax state. The relative shift of the molecular chain of alginate to the arcuate β-turn catalytic channel occurs at the snapshot 00 (12) and snapshot 06 of the lowest energies of alginate, and then the next cleavage triosaccharide oligosaccharide reaction occurs.

**Figure 3.**
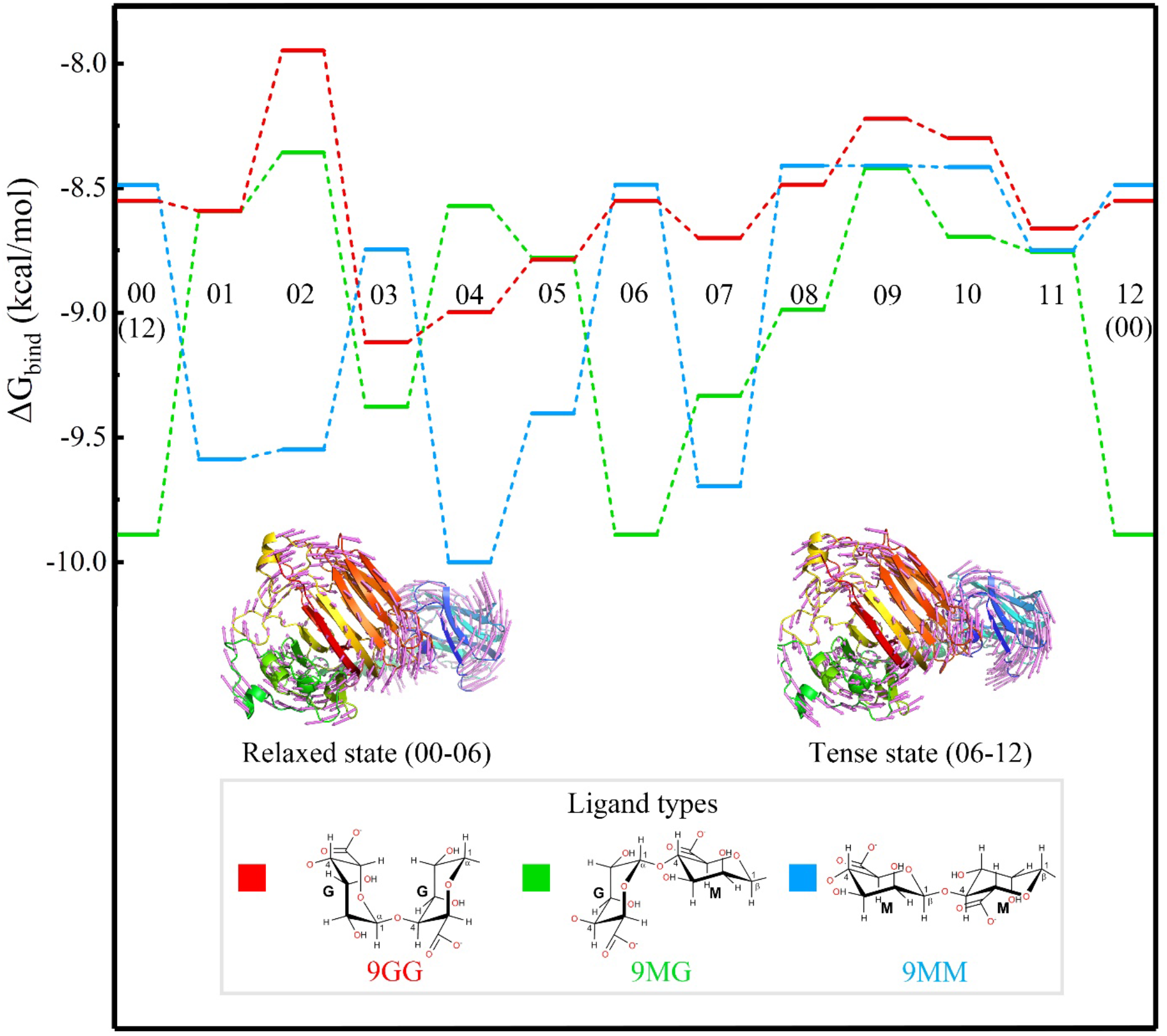
Normal mode-docking analysis of alginate (Nine-mer polyM).

In the energy analysis of Normal mode-docking, we observe that the conformation-docking energy of natural alginate docking to AlyB varies regularly in time (binding energy first rise and then fall in snapshots 00-06 and 06-12). This regular energy cycle variation may be potentially related to the precise cleavaging of natural alginate by AlyB. In the energy change of artificial alginate 9MM (blue), the lowest conformation-docking energy was observed at snapshot 05, indicating that AlyB is most easily bound to 9MM at snapshot 05. Unlike natural alginates, we found that their conformation-docking energy varies irregularly. The conformation-docking energy diagrams of artificial alginate 9MM (blue) and natural alginate 9MG (green) are quite different. As for artificial alginate 9GG, its conformation-docking energy with AlyB (red) is generally higher than that of 9MG and 9MM, indicating that 9GG is more difficult to bind to AlyB than 9MM and 9MG. In addition, in the artificial alginate 9GG, the periodic change of conformation-docking energy similar to that in the natural alginate-AlyB system was not observed.

### CBM32 domain plays the role of grabbing trisaccharide oligosaccharides in degradation mechanism of AlyB

In order to further understand the role of the AlyB subdomain in precise cleavaging biodegradation, we conducted 200ns kinetics simulations of AlyB, AlyB and 9MG, respectively. It was observed that the RMSD of pure AlyB fluctuated little between 0ns-100ns, with a sharp rise around 120ns and gradually leveling off after 150ns (Figure S4). We did not observe a sharp increase in RMSD at 120ns in the kinetic simulation of AlyB and 9MG (Fig. S5), which indicated that the combination of alginate and AlyB was conducive to the stability of the whole system. In addition, we observed that RMSF of 0-150 residues in both AlyB and AlyB-9MG was significantly higher than that of other residues, which were all located in the CBM32 domain, indicating that the CBM32 domain was very flexible relative to other parts of AlyB.

Then, in order to further explore the direction of triosaccharide oligosaccharide products, we conducted a 200ns kinetics simulation with AlyB and 3MG. In the simulation of AlyB-3mg, we observed that the RMSD of the system was basically stable after 75ns. Six representative structures, a, b, c, d, e, f were intercepted in the whole 200ns simulation (Figure 4a, 4c). After superposition, it was found that the positions of the CRM32 domain and Helix-linker changed greatly in the whole simulation, while the positions of the PL-7 domain changed slightly. This suggests that the CBM32 domain performs a different biological function than the PL-7 domain. More interestingly, as shown in Figure 4b and 4d, we compared the RMSF of pure AlyB system with that of AlyB-3MG system, and found that the addition of 3MG increased the RMSF of the R1, R2 and R3 regions of AlyB. While the R1 region is located in the CBM32 domain, the R2 region is corresponding to the Helix-linker and CD-linker, and the R3 region is located in the PD-7 domain. The increase of RMSF in the residues of these structural regions also suggests that they play a direct or indirect role in the release of triosaccharide oligosaccharide products.

**Figure 4.**
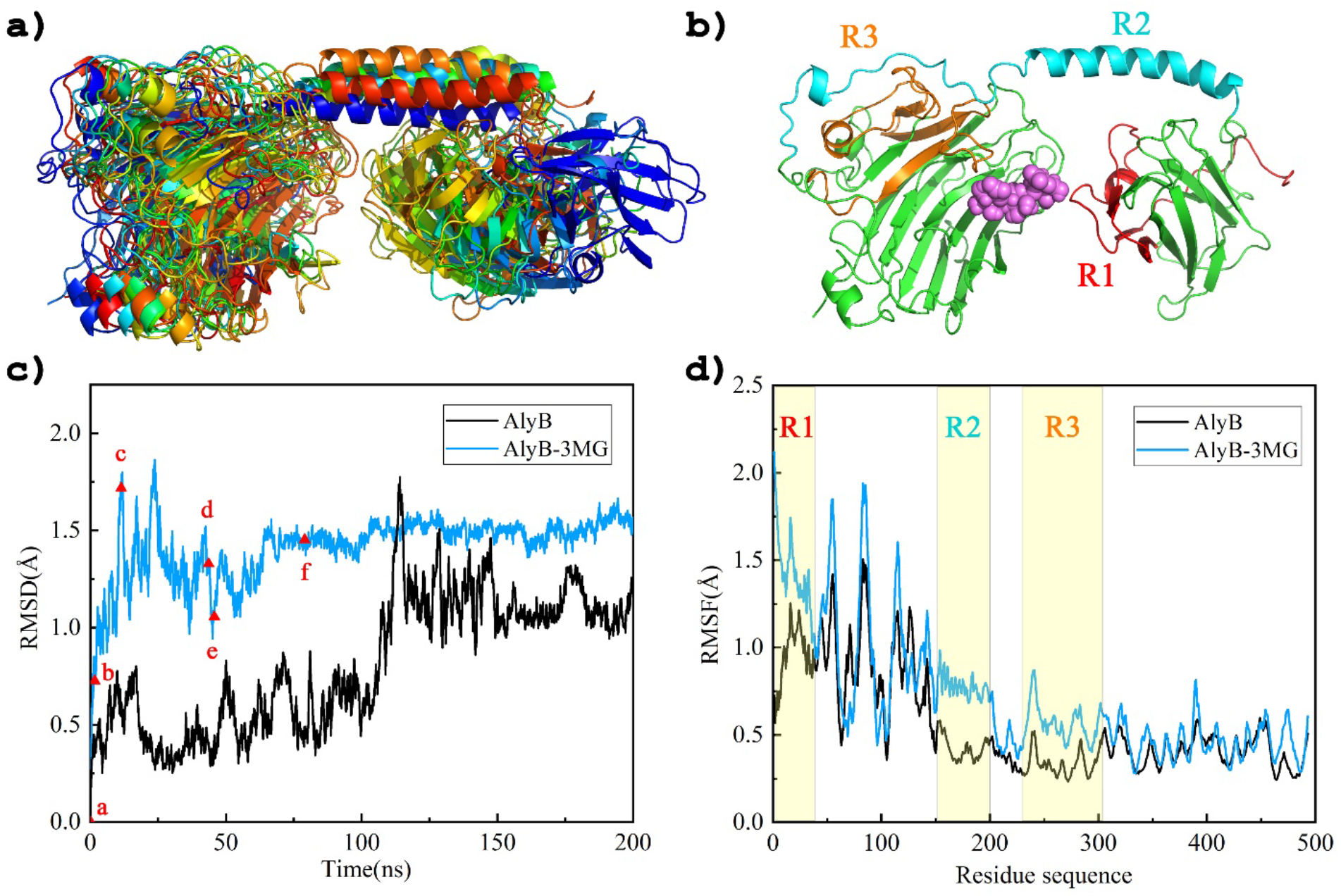
The RMSD and RMSF of Kinetic process of capturing trisaccharide oligosaccharides by CBM32 domain.

In order to more clearly and visually understand the direction of triosaccharide oligosaccharides, six representative structures a, b, c, d, e, f of the kinetic system of AlyB-3MG were demonstrated in fig.5. Fig. 5a is the initial structure of the AlyB-3MG system, which is located at the right end of the arcuate β-turn catalytic channel, and is the initial location of the β-elimination cleavage to produce 3MG triosaccharide oligosaccharide in the AlyB-9MG system. From Fig. 5a to Fig. 5b, we observed that the CBM32 domain was far away from the PL-7 domain, while 3MG rotated to some extent, but the spatial position changed little. In Fig. 5b to Fig. 5c, we found that the CBM32 domain was still far away from the PL-7 domain, but 3MG moved towards the Helix-linker direction. In Fig. 5c to Fig. 5d, the distance between the CBM32 domain and the PL-7 domain gradually narrowed, but there was almost no change in 3MG. Finally, from Fig. 5d to Fig. 5e and then to Fig. 5f, the CBM32 domain approaches the PL-7 domain and finally binds to 3MG. The whole dynamic process of capturing the trisaccharide oligosaccharide 3MG was completed by the CBM32 domain.

**Figure 5.**
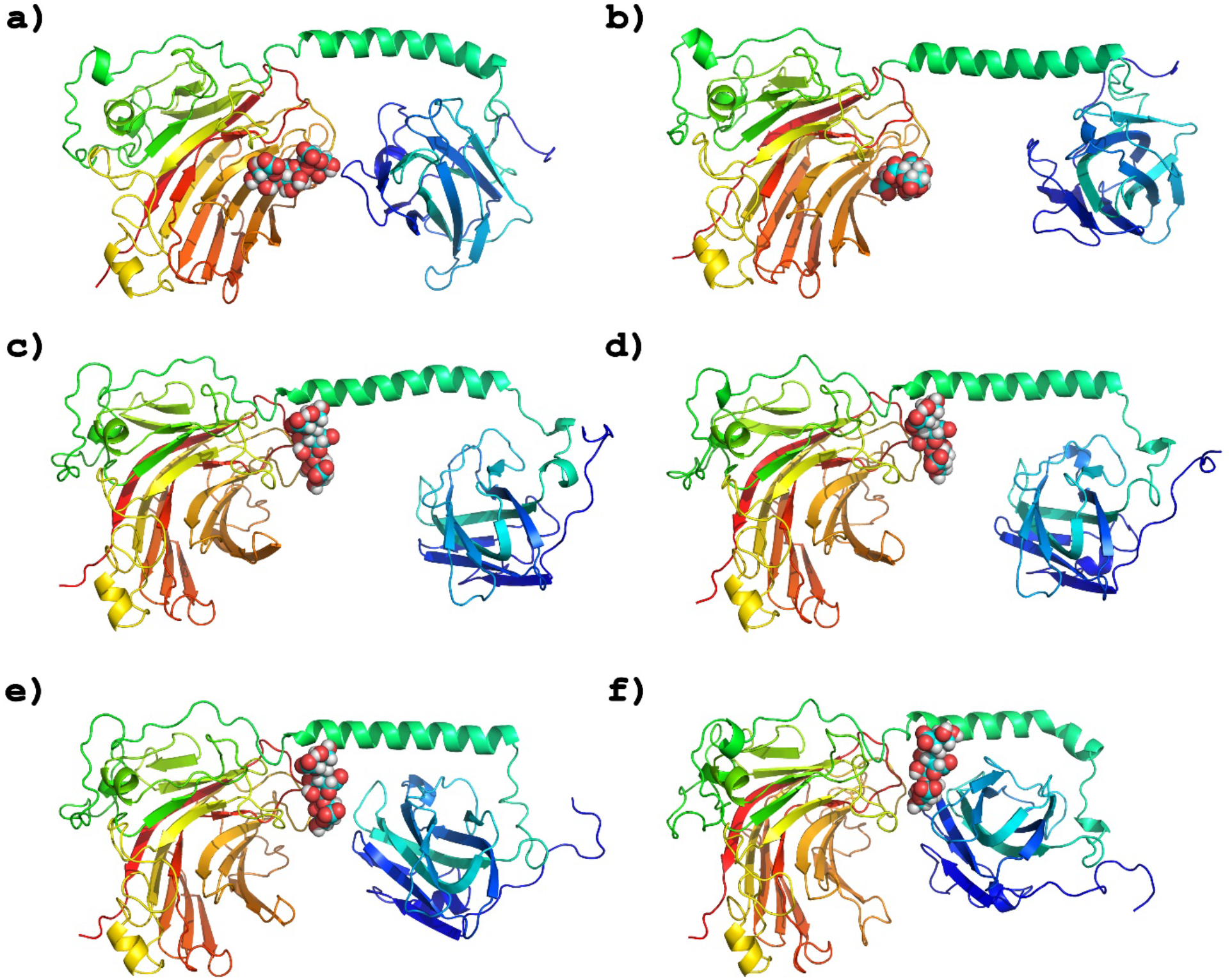
Kinetic process of capturing trisaccharide oligosaccharides by CBM32 domain.

**Figure. 6.**
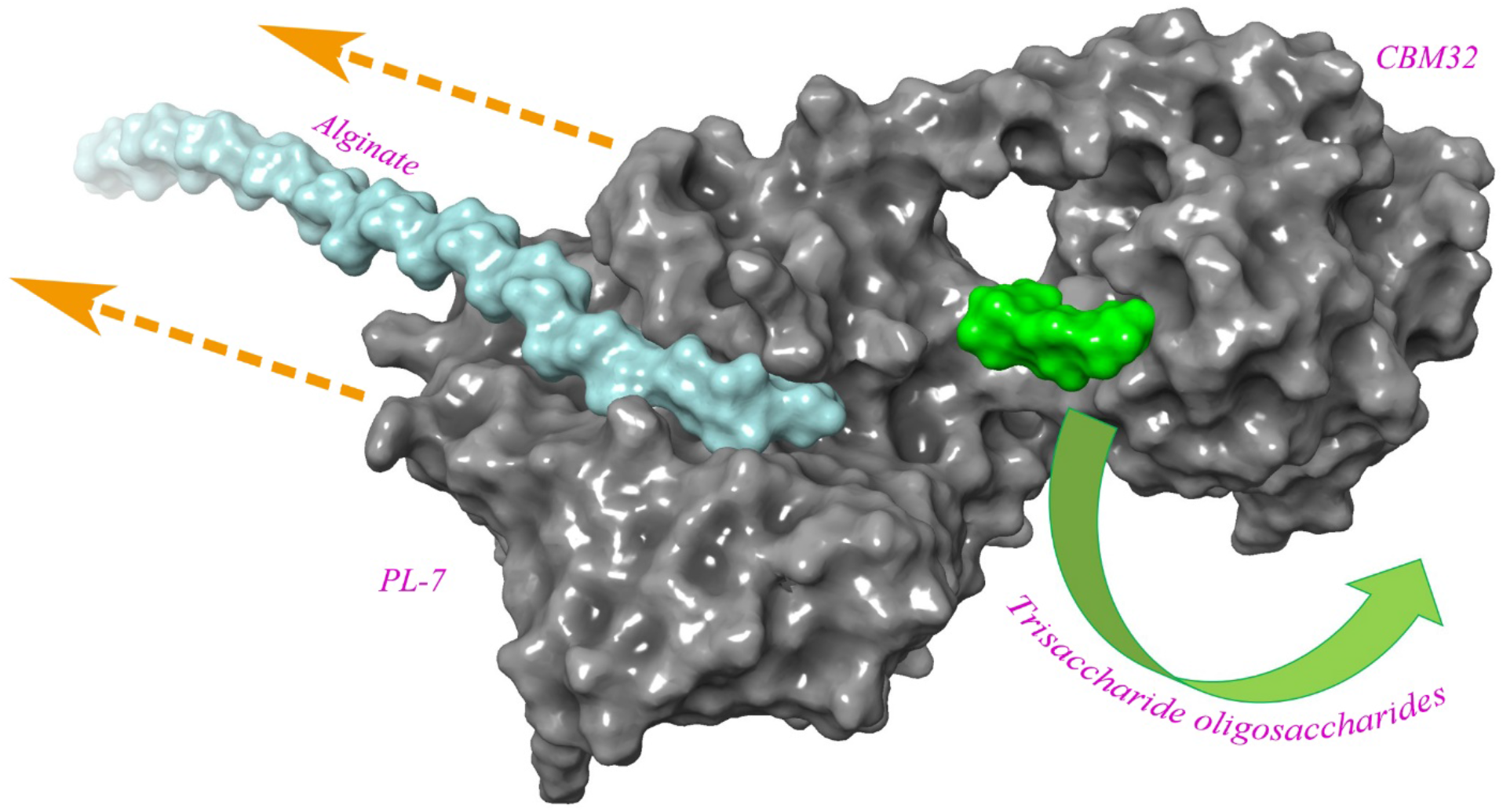
The precise cleavaging bio-enzymatic degradation mechanism of AlyB. (The alginate chain is blue, the trisaccharide is green)

### The precise cleavaging bio-enzymatic degradation mechanism of AlyB

Based on the above research results, we believe that the molecular mechanism of cleavage of alginate by lyase may be as follows: The lyase moves along the long chain of alginate components, and its own structure is constantly twisted to form periodic energy fluctuations. During the relax state of AlyB, the alginate molecular chain enters the arcuate β-tune catalytic channel from the left end of PL-7, and then AlyB transforms from relax state to tense state, and the trisaccharide oligosaccharide is precisely cleavaged by β-elimination reaction. In quick succession, AlyB changed from tense state to relax state, and CBM32 grabbed the trisaccharides oligosaccharides. AlyB then moved in the orange arrow direction (in Fig.6), and the alginate molecular chain shifted three sugar rings to the right relative to AlyB, while CBM32 released the triosaccharide oligosaccharide. Finally, AlyB is transformed from a relax state to the tense state again to carry out the next precise cleaving catalytic cycle.

## Conclusions

As an important polysaccharide natural resource, alginate is a very important raw material for biological manufacturing and pharmaceutical industry. Its production depends on specific alginate lyase. As a natural alginate lyase, most of its products are either monosaccharides or mixtures of a variety of oligosaccharides. It is very difficult to obtain mono-distributed pure oligosaccharides by separating these mixtures, which greatly limits the industrial application of these enzymes. Among various alginate lyases, those that can degrade specific types of alginate substrates and produce quasi monodisperse oligosaccharides have attracted special attention due to their potential bioengineering and medical applications.

In order to investigate the substrate specificity and product uniformity of alginate lyase, we studied the degradation mechanism of a multi-domain alginate lyase AlyB isolated from *Vibrio spendidus OU02*,an enzyme that degrades the natural alginate to produce mono-distributed tri-saccharides. We compared the interaction models between the enzyme and different polysaccharide substrates. Through docking analysis based on normal mode and molecular dynamics simulation calculation, we quantitatively gave the potential mechanism of its cleavage of natural alginate to produce tri-saccharides. Our calculations show that the catalytic pocket of AlyB spans six sugar units, and the loose arrangement of the two domains inside the complex structure endows the enzyme very flexible conformational changes. This arrangement makes the enzyme bind more tightly to natural sodium alginate (PolyMG) than to artificially designed polysaccharides (PolyM and PolyG). At the same time, under this favorite binding conformation, the spatial orientation of the catalytic site is conducive to the cleavage of the glycosidic bond between the third and fourth sugar units, resulting in single tri-saccharide products. On the other hand, molecular dynamics simulations showed that the C-terminal CBM domain can search a wide range of orientations related to the catlytilic domain, thus being able to capture the cleaved tri-saccharide product from the catalytic domain, and taking them away timely from the catalytic center through an inherent motion by itself, so as to avoid further hydrolysis of the products to produce monosaccharides or disaccharides. Our calculations suggested that the linkage of AlyB and CBM domains through the long helix structure endows the enzyme a critically important capability of conformation changes, which plays a decisive role in the specific selection of substrate, and determines the uniformity of the products. Our data provided new insights for the rational design of new enzymes that may contribute to the production of monodisperse oligosaccharides.

## Acknowledgements

This work was supported, in part, by the National Key Research and Development Program of China (Grant No. 2019YFA0905700, 2017YFC1600900). Part of the work was carried out when one of the author (H.X.) visited Tianjin Institute of Pharmaceutical Research.

