## Supplementary figures and images for "How alginate lyase produces quasi-monodisperse oligosaccharides: a normal mode analysis-based docking and molecular dynamics simulation study"

### Supplemental Figure S1

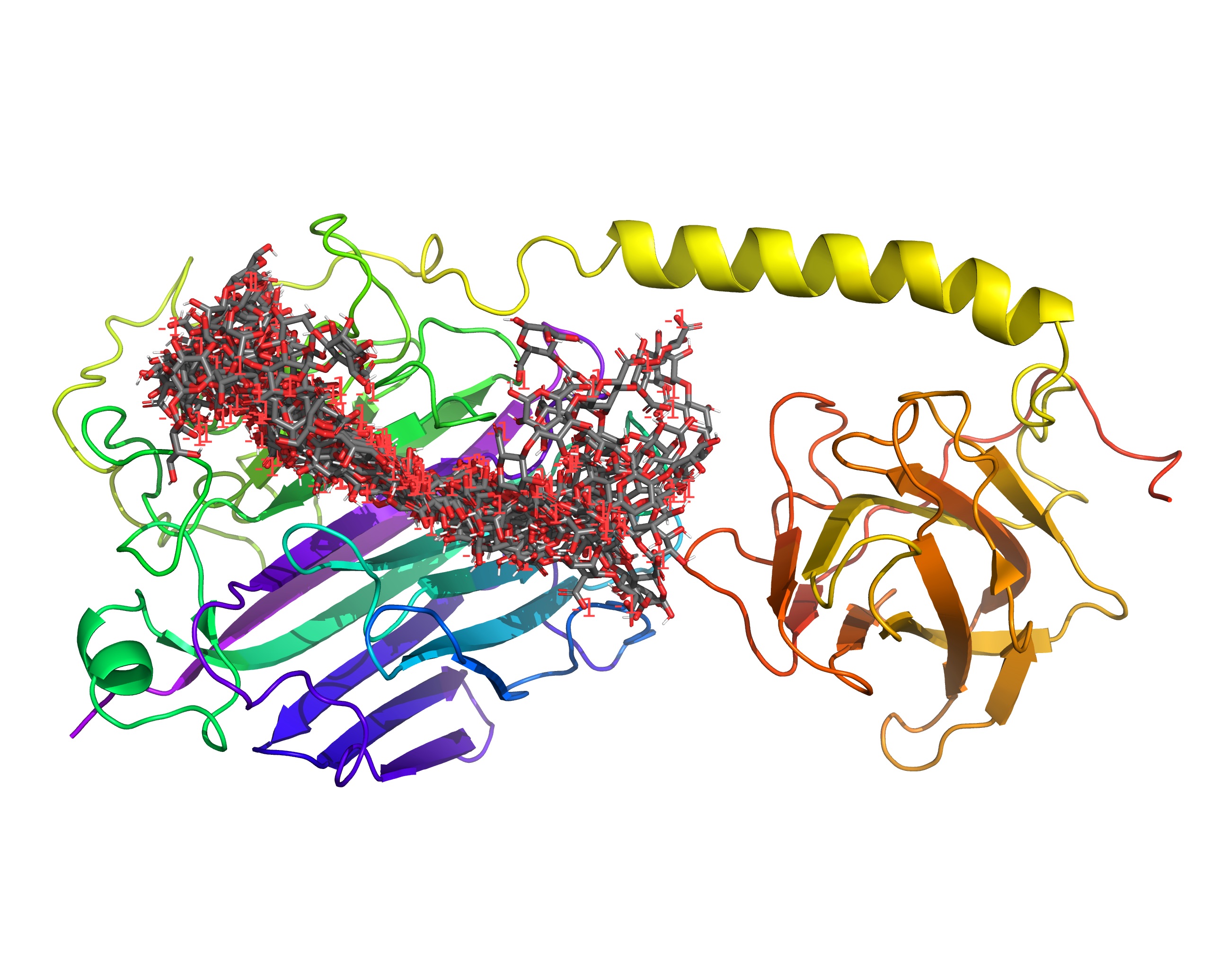

### Supplemental Figure S2

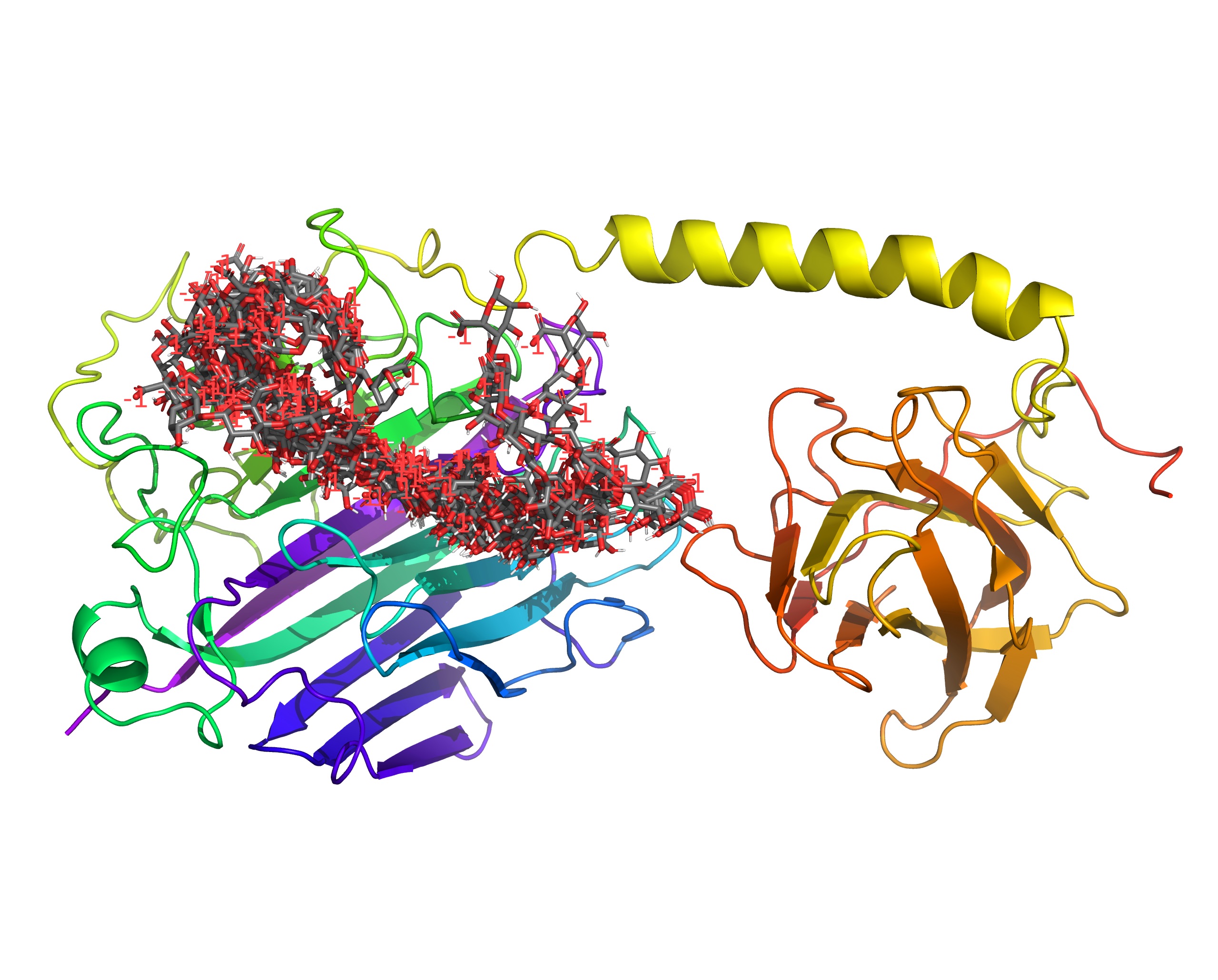

### Supplemental Figure S3

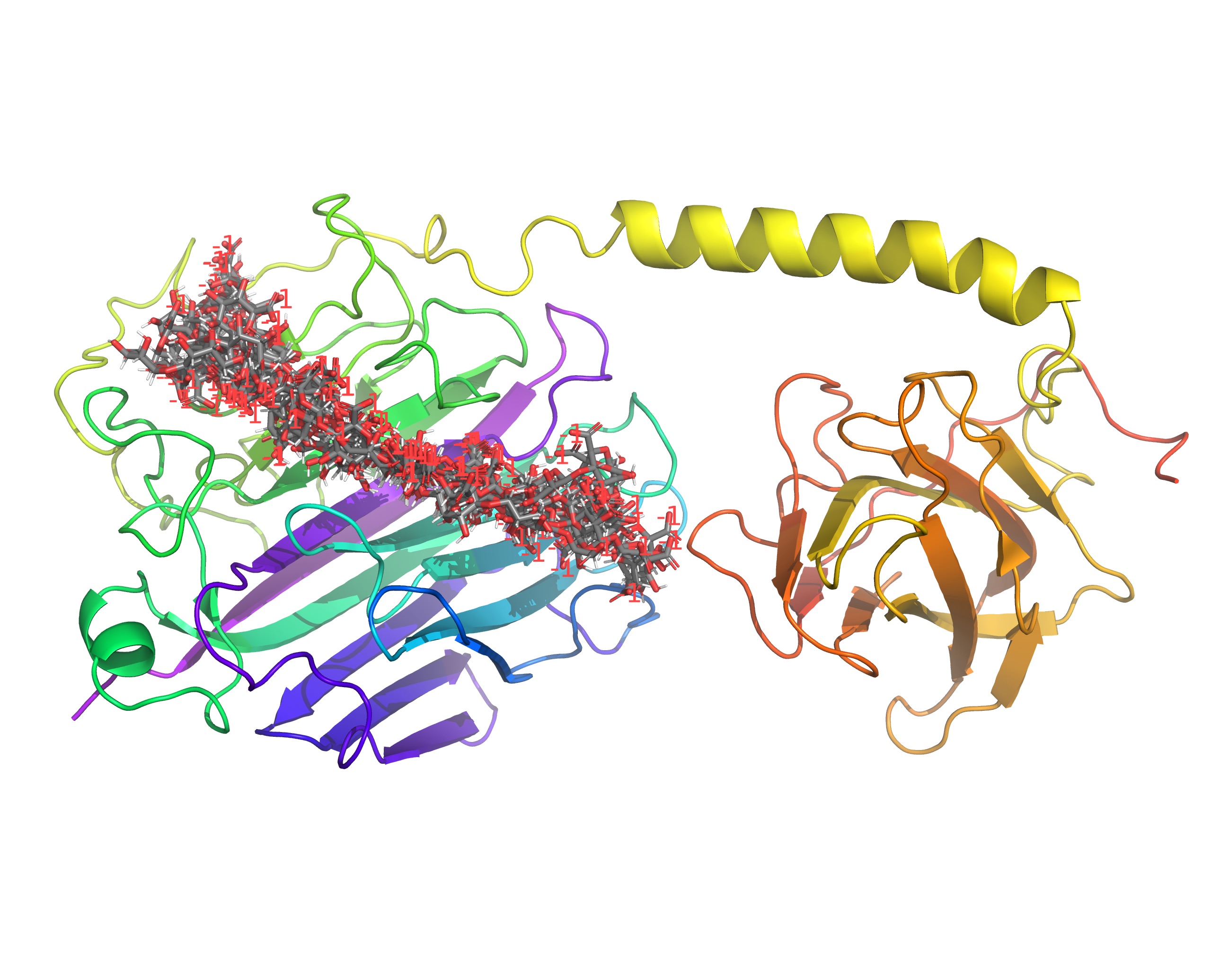

### Supplemental Figure S4

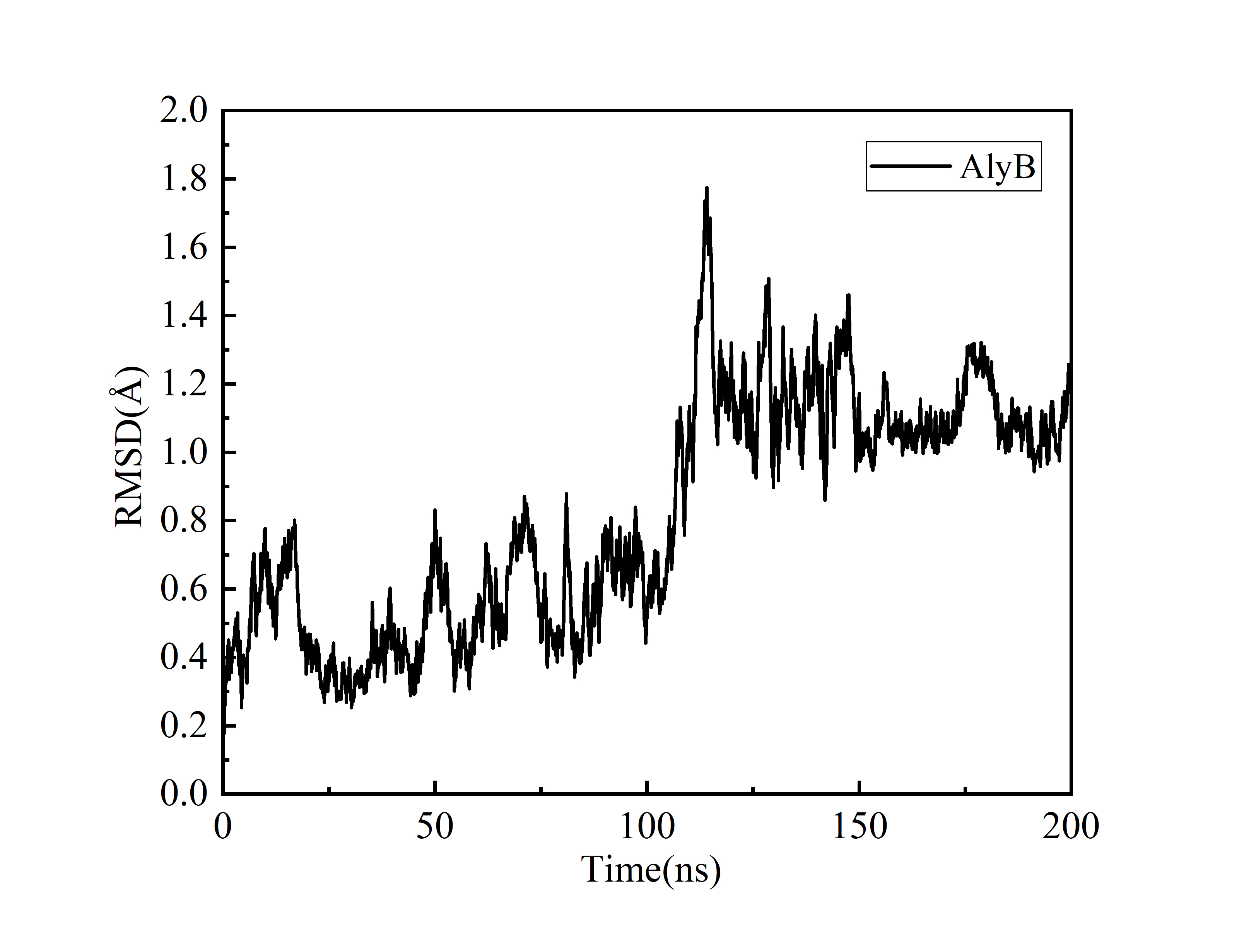

### Supplemental Figure S5

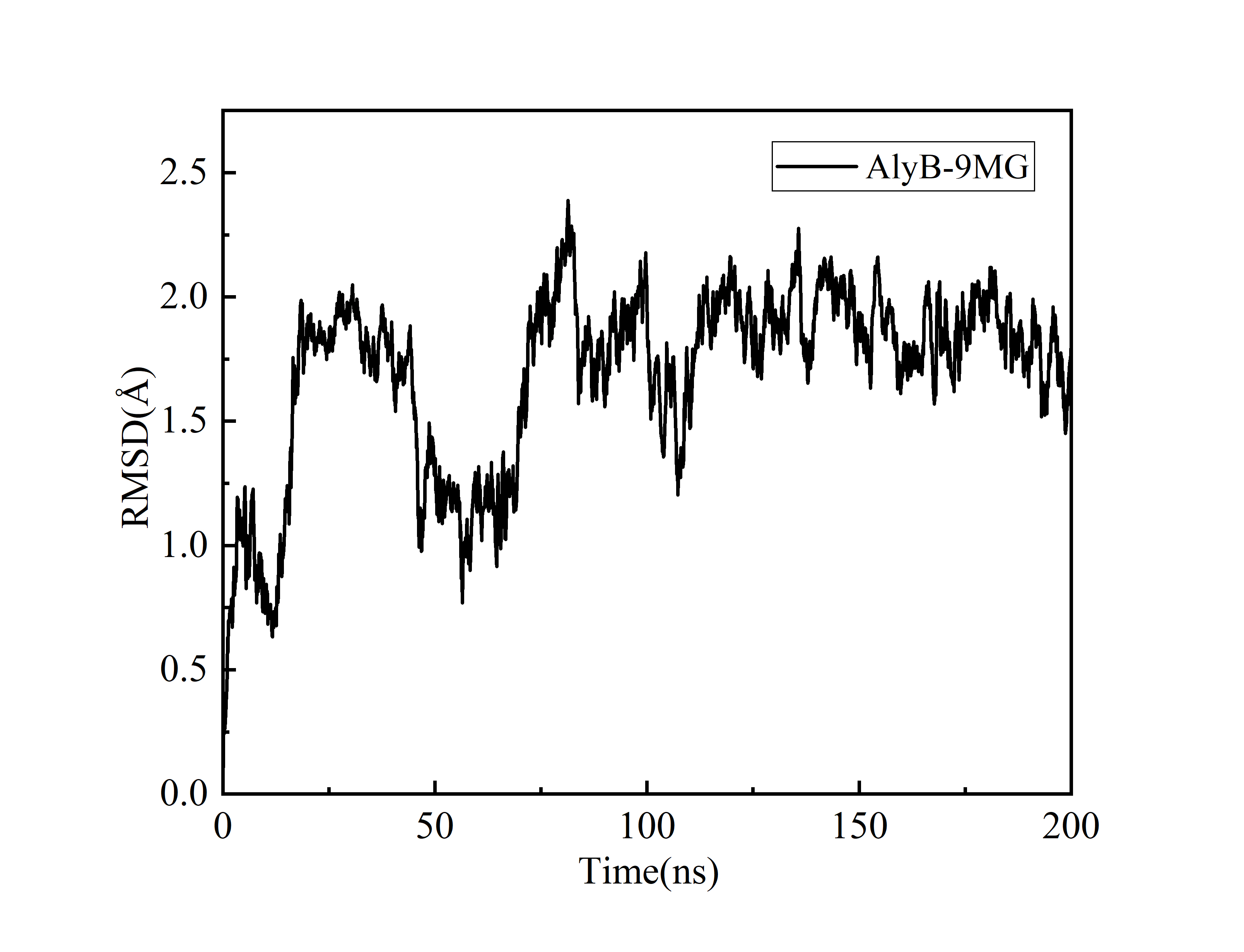

### Supplemental Figure S6

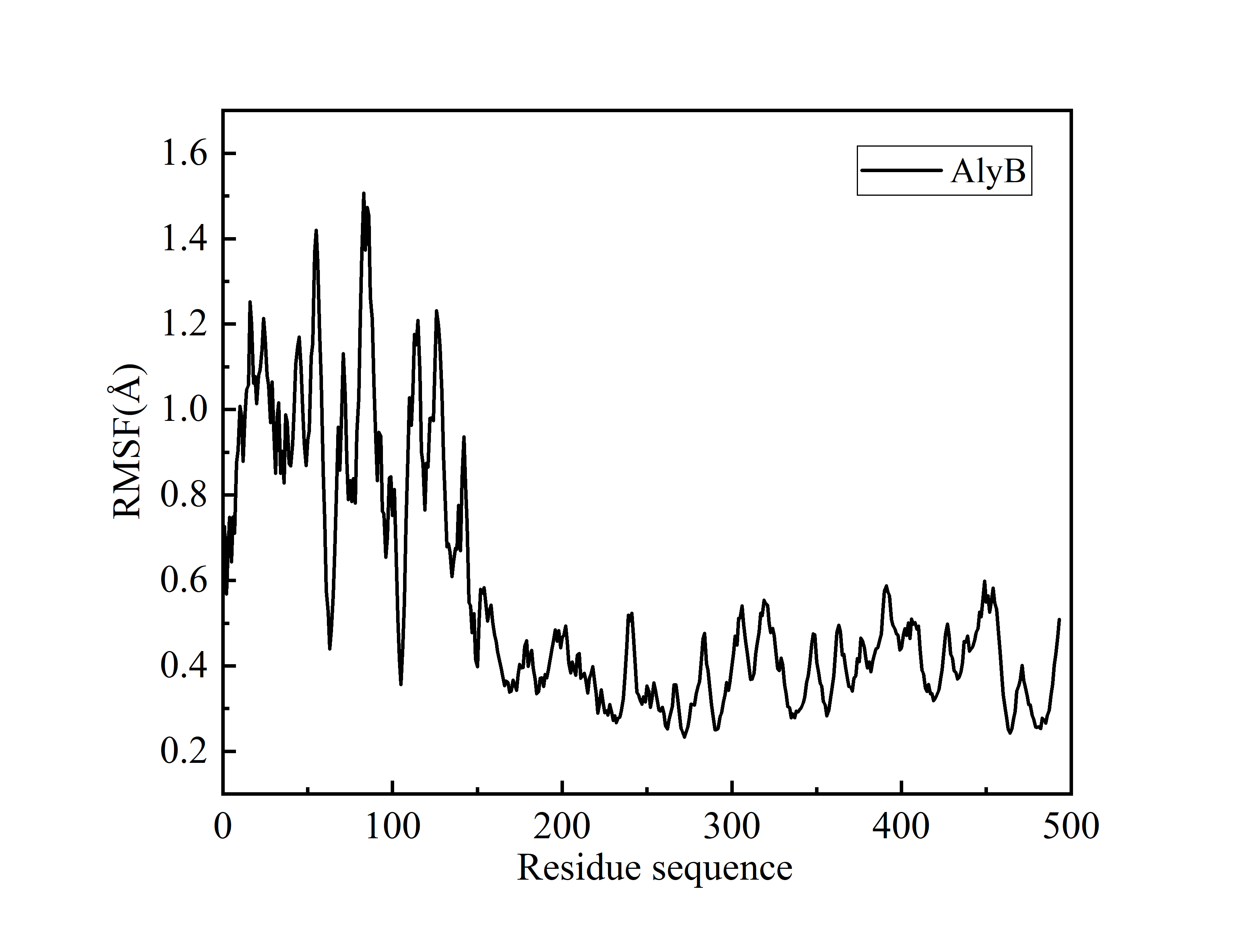

### Supplemental Figure S7

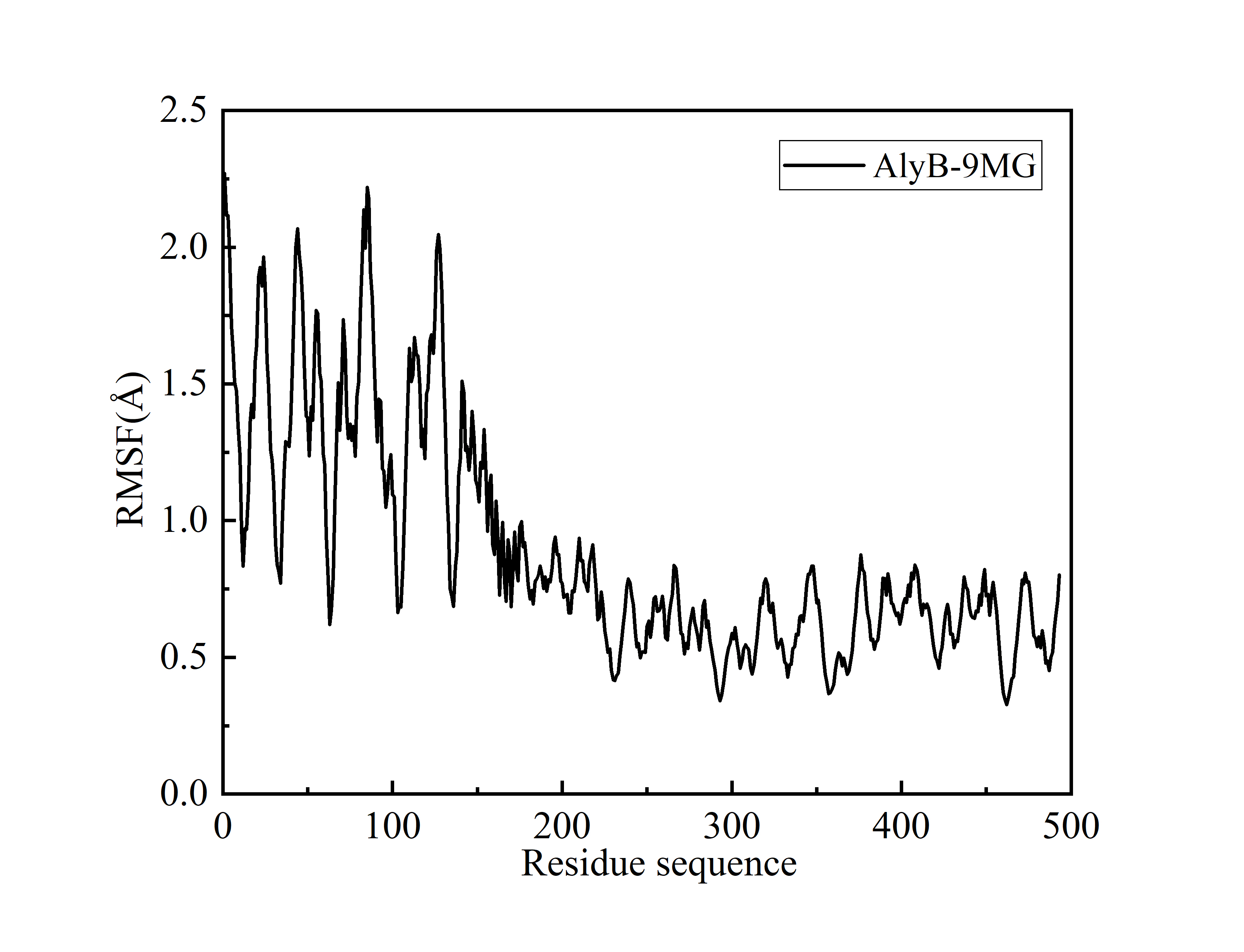
